# Inversed association of locus coeruleus MRI integrity with structural volume and its impact on individual’s inattentiveness

**DOI:** 10.1101/2024.09.27.614628

**Authors:** Joshua Neal, Sun Hyung Kim, Benjamin Katz, Il Hwan Kim, Tae-Ho Lee

**Affiliations:** Department of Psychology, Virginia Tech, USA; Department of Psychiatry, University of North Carolina, Chapel Hill, USA; Department of Human Development and Family Science, Virginia Tech, USA; School of Neuroscience, Virginia Tech, USA; Department of Anatomy and Neurobiology, University of Tennessee Health Science Center, USA

**Keywords:** Locus coeruleus, MRI contrast, integrity, structural volume, inattentiveness, attention

## Abstract

The locus coeruleus (LC) is a nucleus within the brainstem associated with physiological arousal and altered structure and function in the context of neurodegenerative disorders. Pathologies related to difficulties with attention have previously been associated with abnormalities in neurotransmitter production and sensitivity, suggesting the possibility of abnormality in neurotransmitter producing neural regions. One such region is the LC, associated with norepinephrine production. To examine the possibility that LC alteration is associated with inattentive symptom reporting, a set of analyses have been performed with 141 individuals age-ranged from 8 to 54. We found that the structural integrity value of the LC, especially on the right hemisphere, showed a significant negative association with the level of individual’s inattentiveness score. Furthermore, LC volume size was significantly positively associated with inattention, and this finding was also lateralized to the right LC. Interestingly, an inverse association was found between structural integrity and volume size. These findings support the relationship between LC and attention-related behavior through both neuromelanin-sensitive and structural imaging, with important implications for the association between regional structure and function.

## Introduction

Attention is vital to our daily life (Carrasco, 2011) as it is the cognitive process of assigning limited mental resources to differing thoughts, concepts, memories, or stimuli at any given time. Significant attentional deficits are thus associated not only with the difficulties in daily activities but also with development of psychopathological symptoms often reported from attention deficit/hyperactivity disorder (ADHD) Though there are findings supporting environmental influences upon the development and severity of inattention (Johnson et al., 2019; C.-Y. Liu et al., 2019; Mikami et al., 2010; Neugebauer et al., 2015), much of the literature regarding these attentional difficulties focuses on neurological abnormalities that may be linked to observed behaviors.

The neurological differences implicated in attention deficits span different neural regions and functions. Altered serotonergic (Quist et al., 2003; Vanicek et al., 2017; Zepf et al., 2010) and noradrenergic (Biederman & Spencer, 1999; Russell et al., 2005; Yang et al., 2013) systems in the brain have been observed in individuals displaying inattentive symptoms, with abnormalities either reducing the production of the given neurotransmitter or limiting their efficiency in synaptic transmission. Medications that reduce inattention symptoms are thought to do so mainly by increasing the effects of these neurotransmitters within the frontal and prefrontal cortex regions (del Campo et al., 2011; Wilens, 2008). Given findings connecting subcortical regions with neurotransmitter production and signaling (e.g., (Alcaro et al., 2007; Berridge & Waterhouse, 2003; de la Cruz et al., 2021; Teissier et al., 2015), neural models of inattention should seek to integrate subcortical regions associated with neurotransmitter production into models of neural attention.

One potential cause of altered neurotransmitter production or signaling is the abnormality or neurodegeneration of the locus coeruleus (LC), a brainstem nucleus recognized as the primary site of norepinephrine production in the brain (Nai-shin & Bloom, 1973; Nieuwenhuis et al., 2005; Redmond & Huang, 1979; Sara, 2009). While the LC has been associated with broad physiological arousal in individuals, accumulated evidence suggests that it could send inputs to various brain regions uniquely related to a task-specific demands. For example, Clewett et al (2018) showed that the LC increases its functional connectivity with the parahippocampal place area (PPA) in processing of memory consolidation of place/scene target images. Similarly, Lee et at (2014) demonstrated that the LC showed strengthened connectivity with the fusiform face area (FFA) when an individual was required to detect fact target stimuli in the selective attention task. Lee et al. (2018) further demonstrated that such differed LC signaling can vary by age factors, with optimal stimuli task performance found for younger individuals exhibiting not just higher selective LC activation, but in higher functional connectivity (FC) with task specific networks (e.g., frontoparietal network, FPN) to process target stimuli depending on the task demands. In contrast, older adults were found to maintain neural arousal often, but the loss of LC-FPN FC was associated with decreased ability in selective processing. Similarly, animal studies demonstrated that LC sends unique neural inputs differently to the sensory network regions in mice (Devilbiss & Waterhouse (2011). These findings suggest the LC may act as a neuromodulatory signal mechanism to cortical networks and may influence the cognitive processes or saliency of a network through differential activation.

However, the LC has been limited in standard structural estimates, primarily because it is a very small structure with an average width and depth of 2.0 mm in the brainstem (Fernandes et al., 2012). Structural estimation such as volume size based on these observations in ex vivo examinations would correspond to 50mm^3^ per hemispheric portion, whereas MRI based estimates have ranged 8-30 mm^3^ across prior studies (Castellanos et al., 2015; X. Chen et al., 2014; Langley et al., 2017; S. T. Schwarz et al., 2017). Recent improved methodologies involving multi-atlas probability estimates (MacDuffie et al., 2020; Shen et al., 2022; Utsumi et al., 2020; Ye et al., 2021) present volumes more in line with anatomical studies, however, there is still difficulty in estimation with evidence of heterogenous projections along the length of the cylindrical nucleus (Ao et al., 2021; Chandler et al., 2014; Plummer et al., 2020). Thus, the structural characteristics of the LC should be accurately examined in conjunction with other measures, such as those providing insight into neurotransmitter production.

Given difficulties with structural estimation, the LC has been assessed instead using a neuromelanin contrast ratio value (CNR)(Clewett et al., 2016; Sasaki et al., 2006). The unique properties of neuromelanin as normally concentrated within the region produce a significantly higher signal intensity value during a structural MRI scan, which can then be computed into a contrast ratio against a reference region to estimate the integrity and function of the LC. The resulting neuromelanin contrast ratio has been associated with cognitive performance (Chowdhury et al., 2022; Clewett et al., 2016; Hämmerer et al., 2018; K. Y. Liu et al., 2020) as well as the onset and progression of neurodegenerative diseases such as Alzheimer’s and Parkinson’s (Olivieri et al., 2019; Putcha et al., 2023; Takahashi et al., 2015; J. Wang et al., 2018). Neuromelanin contrast has also been found to be associated with other structural values, such as cortical thickness, volume, and microstructure integrity (Bachman et al., 2021; Elman et al., 2022; Taniguchi et al., 2018). Asymmetries in neuromelanin contrast values between hemispheres have been previously found and connected to symptom presentation (Kuya et al., 2016; Shinde et al., 2019; Trujillo et al., 2023; J. Wang et al., 2018). In particular, within Parkinson’s disease lateralized differences in CNR were connected to differences in motor degeneration by hemisphere of the body (Prasad et al., 2018), establishing that asymmetry may be a representative measure of qualities pertaining to neuromelanin hemispheres and associated psychophysiological effects. A separate study (Q. Liu et al., 2023) established lateralized differences in LC CNR for Parkinson’s patients, and associated those values to non-motor related symptoms. To date, there have not been significant studies of LC neuromelanin contrast in neurodevelopmental disorders, but existing literature has identified associations between norepinephrine with ADHD and ASD (Angyal et al., 2018; C.-H. Kim et al., 2006; Y. Kim et al., 2022; Launay et al., 1987), suggesting neuromelanin contrasts for noradrenergic-producing regions such as the LC may be associated with attention processing outside of a neurodegenerative context.

As such, the present study has further evaluated the potential of LC attributes being related to the presence of inattentive symptoms. The overall aim was to examine structural and metabolic values for the LC in a general population sample, examining the possibility that these values may explain attention behaviors and difficulties outside of neurodegenerative conditions. As neuromelanin is understood to be positively related to norepinephrine production, we hypothesized that greater concentrations of neuromelanin would be associated with greater attentive behavior, or that lower neuromelanin levels would be associated with higher report of inattentive behaviors.

Consistent with this aim, we conducted multiple analyses with structural and neuromelanin brain images based on a typically developing (TD) population. Behavioral reports of inattentive behaviors were collected, as were standard and neuromelanin-sensitive structural images. Each hemisphere of the LC was examined separately in relation to inattentive behavior reporting.

## Method

### Participant characteristics

The 141 healthy participants were recruited and scanned as part of larger lab’s cross-sectional brain imaging project (*M_age_* = 25.04; *SD* = 13.53; range = 8 - 54; female = 51.77%; see Table 1 for demographic information). All participants had normal or corrected-to-normal vision and hearing, and self-reported no history of chronic illness or cognitive impairment. 19 participants reported the use of prescription medication at the time of the scan including fluvoxamine, sertraline, welbutrin, and lexapro, and the medication status was controlled in the analysis. All participants provided written informed consent approved by Virginia Tech Institutional Review Board.

**Table 1.** Descriptive statistics of age, sex, and race for the study population.

| Variable | Age (in years) | Sex | Race/ethnicity |
| --- | --- | --- | --- |
| Mean (M) | 25.04 |  |  |
| SD | 13.53 |  |  |
| Range | 8-54 |  |  |
| Male |  | 68 (48.23%) |  |
| Female |  | 73 (51.77%) |  |
| White |  |  | 98 (68.09%) |
| Black |  |  | 3 (2.13%) |
| Asian |  |  | 36 (25.53%) |
| Other |  |  | 4 (2.84%) |

### MRI data acquisition

To examine the relationship between LC and attention deficits, T1w/T2w structural images and neuromelanin-sensitivity structural images were collected with 3T Siemens MRI scanners at the FBRI Biomedical Research Institute at Virginia Tech Carilion (FBRI-at-VTC) and the Virginia Tech Corporate Research Center (VTCRC) respectively. We included this information as a nuisance regressor in the analyses to control for potential scanner and location variabilities. One neuromelanin-sensitive-weighted MRI scan was collected using a T1-weighted FSE imaging sequence (repetition time = 750 ms, echo time = 12 ms, flip angle = 120°, 1 average to increase signal-to-noise ratio (SNR), 11 axial slices, field of view = 220 mm, bandwidth = 285 Hz/Px, slice thickness = 2.5 mm, in-plane resolution = 0.43 × 0.43 mm). We also collected a high-resolution anatomical image for each participant (T1-weighted image: repetition time = 3200 ms, echo time = 2.06 ms, flip angle = 8°, bandwidth = 220 Hz/Px, voxel resolution = 1 mm^3^ isotropic; T2-weighted image: repetition time = 2,500 ms, echo time =56 ms, flip angle = 120°, bandwidth = 725 Hz/Px, voxel resolution = 1 mm^3^ isotropic).

### Measurement of attentional behavior

To evaluate individual’s attentional behavior, we utilized two attention scales, the *Barkley Adult ADHD Rating Scale IV (BAARS-IV)* and the *Vanderbilt Parental Rating Scale for children.* These two scales are primarily used for clinical diagnosis purpose, but scores from these measures are commonly considered as a reliable attention measure for non-clinical samples (Ebert, 2017; L. A. Miller et al., 2015; Pritchard et al., 2017; Wood et al., 2021). In studies of attention behavior across the lifespan, these two scales have been used similarly to evaluate attentional constructs for children and adults (Martel et al., 2012; Marx et al., 2010; Roy et al., 2016), and uniform measure of associated attention behaviors across both children and adults has been validated (Herbert, 2019; Murray et al., 2024). Thus, we used scores from these measures as the quality of individual’s attention behavior.

In measuring attentional behavior, various models have been proposed for the relationships between attentional features in childhood and adults. For example, two factor model primarily considers *inattentiveness* and *combined hyperactivity/impulsivity* in child population (American Psychiatric Association, 1998, 2013)(Gomez & Stavropoulos, 2021; Park et al., 2018). However emerging research suggests that a three-factor model, which distinguishes between *inattentiveness, hyperactivity*, and *impulsivity*, provides a stronger fit for both child and adult populations (Nichols et al., 2017; Parke et al., 2015; Toplak et al., 2012). Given the cross-sectional sample in the current study, which ranged in age from 8 to 54 years, we adopted the three-factor model, utilizing three subscores (*inattentiveness, hyperactivity*, and *impulsivity*) uniformly across two measurements, along with a total summary score.

#### Barkley Adult ADHD Rating Scale IV (BAARS-IV)

BAARS-IV is a self-report rating scale consisting of 18 items corresponding to the nine symptoms of inattention and hyperactivity/impulsivity associated with ADHD (Barkley, 2011). The overall present report value (i.e., total attention score; Cronbach’s alpha = 0.914), as well as the subscales, has been found to have high internal consistency (Cronbach’s alpha for inattention =.902, hyperactivity=.776, and impulsivity=.807) and the test-retest reliability (total attention score =.75, current inattention = .66, current hyperactivity =.72, current impulsivity =.76), as well as inter-rater reliability (Barkley et al., 2011). The present sample also has high internal consistency (Cronbach’s alpha for current total score =.903, inattention =.893, hyperactivity=.724, impulsivity=.821). Each item is evaluated using a 4 item Likert scale, reflecting the occurrence of a behavior within the past six months. Higher score indicates poorer attentional behaviors.

#### Vanderbilt Parental Rating Scale for children

For children and adolescents aged 8-17, parents completed the Vanderbilt Parental Rating Scale (Wolraich, 2003), which measures attentional behaviors based on the DSM diagnostic criteria of ADHD. Reporting consists of 18 items utilizing a 4-level Likert scale, where responses of 3 or 4 were considered indicative of positive report of the associated ADHD symptom within the past six months. Parental report through the measure has previously demonstrated high internal consistency (Cronbach’s alpha for total score =.93), and concurrent validity (r=.79) for parental report total score with structured interview; (Wolraich, 2003). For the present sample high internal consistency was also observed (Cronbach’s alpha = .92).

### Neuromelanin contrast ratio analysis

Neuromelanin contrast was manually collected based on previously established methods (Clewett et al., 2016). For each individual, a T1 FSE image, previously pre-processed and registered to standard space, was opened within FSLeyes (Jenkinson et al., 2012). Within the axial viewpoint (see figure 1) the inferior colliculus was identified, then roughly two slices inferior, the left and right hemisphere LC regions were found, establishing a 3x3 voxel cross surrounding the peak intensity voxel for each LC hemisphere. A 10x10 voxel sample from the dorsal pontine tegmentum, defined as six voxels anterior to the LC crosses and equidistant between them was collected as the averaged reference region for calculating the contrast ratio. An additional set of LC region samples was established one voxel slice inferior to the initial LC region samples. LC’s neuromelanin contrast ratios (LC CNR) are to be calculated for each hemisphere of the LC on the respective axial slices (see equation 1) in addition to a bilateral neuromelanin value. Hemispheric contrasts were also used to calculate neuromelanin contrast asymmetry (see equation 2).

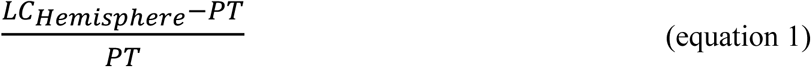

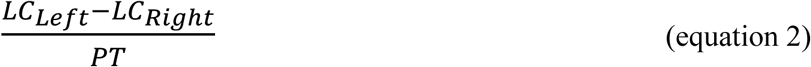

**Figure 1.**
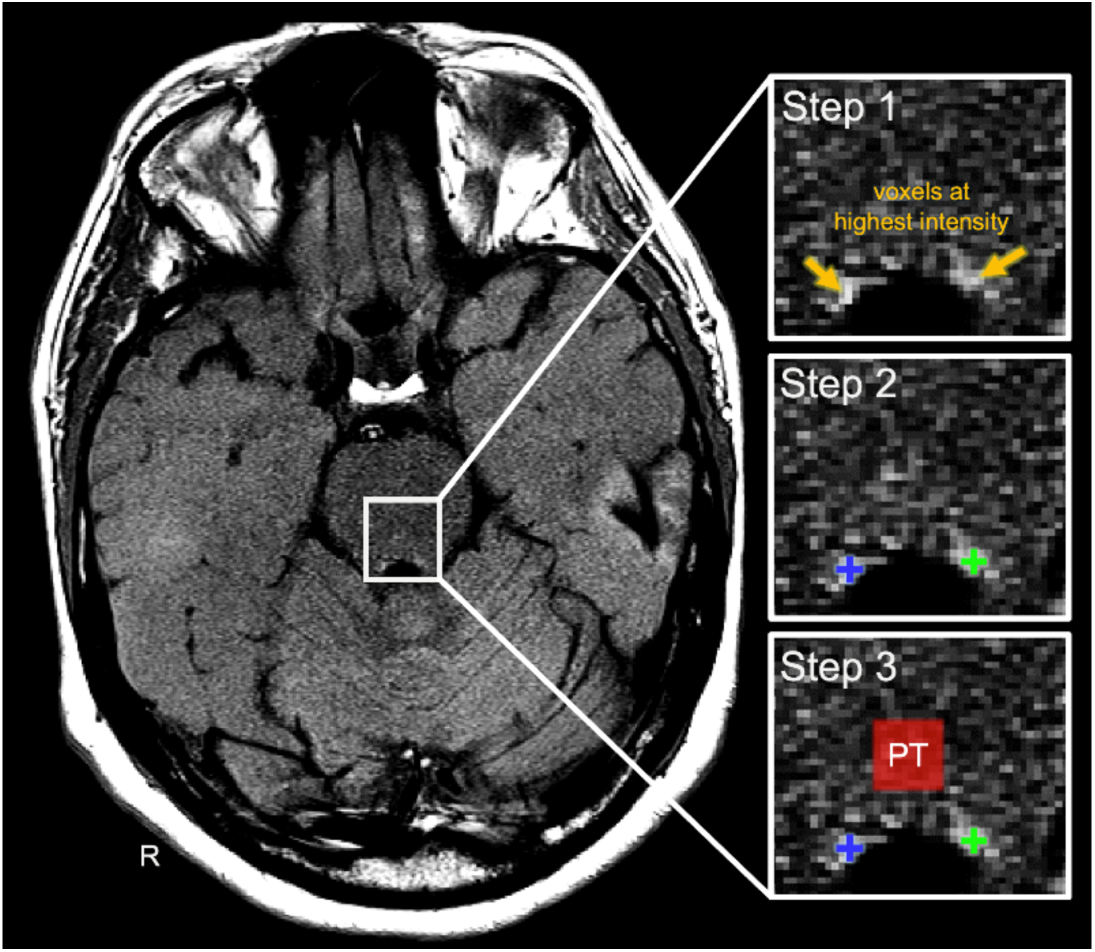
Schematic figure for anatomic tracing protocol. On the axial view of neuromelanin-sensitive MRI image. Step1: Locations of Left and right LC identified as highest signal intensities. Step 2: a small cross (3 × 3 voxels, yellow) of approximately the width of the LC (∼1–2 mm) was placed on the voxels with peak signal intensity. Step 3: The reference, pontine tegmentum (PT), was defined as a 10 x 10 voxel square located 6 voxels above the more ventral of the 2 LCs and equidistantly between them.

## Multi atlas-based LC volume estimation

Quantification of the LC volume was achieved through the integration of neuromelanin-enhanced contrasts delineated manually and the implementation of automated contrast delineation masks.

Prior to estimating automated LC, T1w and T2w brain images were corrected for intensity non-uniformity using the N4 bias field correction algorithm (Tustison et al., 2010). Following this correction, the images were rigidly transformed and aligned with a standardized stereotaxic space. Brain masks were generated through a majority voting approach, which involved the joint warping registration of T1/T2w images to seven predefined single atlases. Manual adjustments to the brain masks were performed using itkSNAP software (Yushkevich et al., 2006) to ensure maximal accuracy.

The locus coeruleus (LC) was defined using a multi-atlas label-based ROI template, complemented by five selected templates from publicly available sources (Edlow et al., 2012; Keren et al., 2009; Liebe et al., 2020; Tona et al., 2017; Ye et al., 2021). A multi-modality (T1w and T2w) multi-atlas segmentation workflow was implemented using the in-house, open-source MultiSegPipeline software (Cherel et al., 2015). This software enabled the application of majority voting techniques following the warping of LC annotations using Advanced Normalization Tools (ANTs) (Tustison et al., 2021). To maintain congruency between the manually delineated LC masks and the automated segmentation outputs, and to rectify any discrepancies in spatial alignment, a morphological operation of one voxel dilation followed by erosion was performed. A thorough visual assessment was performed to evaluate the anatomical accuracy of the segmentations across all images, confirming that no processed data needed to be excluded due to suboptimal segmentation quality. The volume was computed by multiplying the total number of voxels in the combined integrated label with the spatial resolution. Study population averages for the structural estimates are listed in table 2.

**Table 2.**
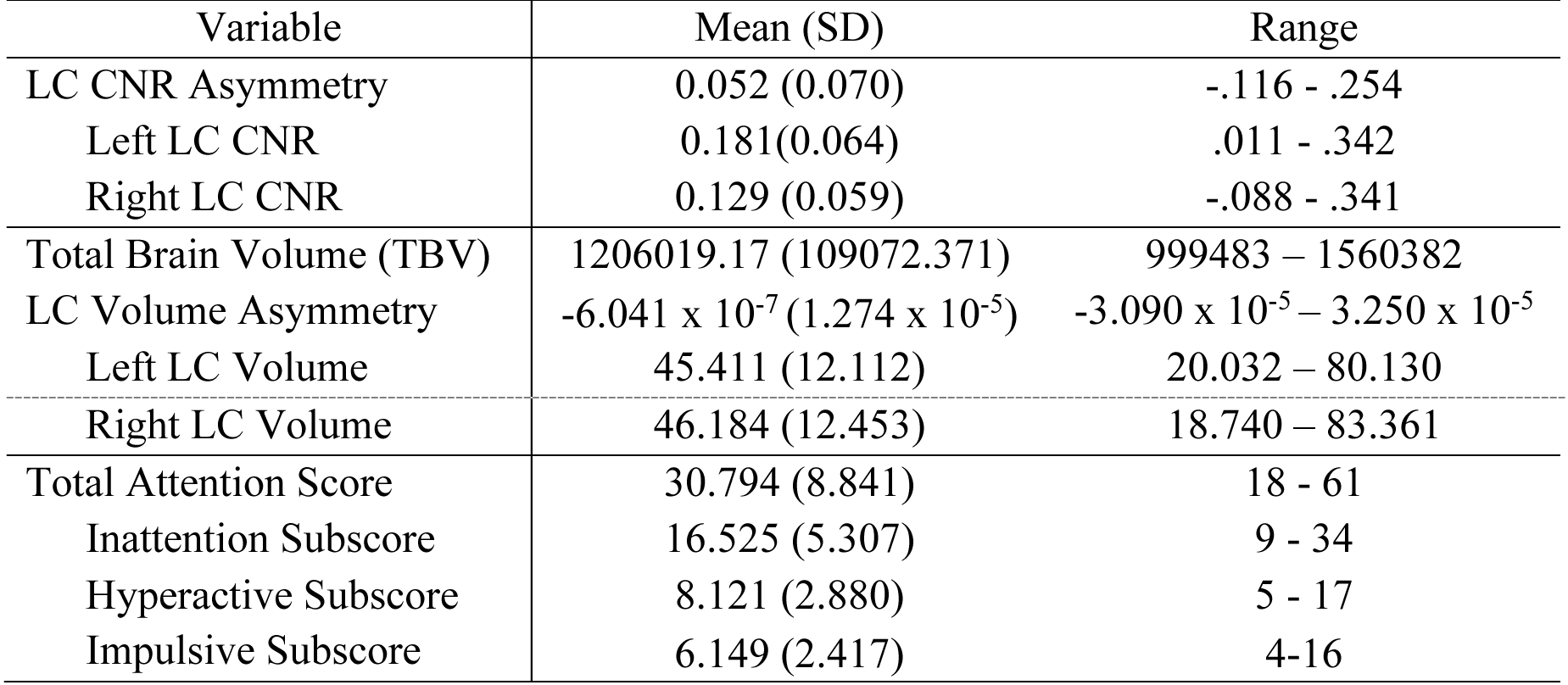
Descriptive statistics of LC CNR, LC volumes and behavioral scores for the study population (SD: Standard deviation; CNR: Contrast-noise-ratio)

## Regression analyses

LC neuromelanin contrast values were regressed separately onto both the score total for the respective behavioral report as well as the number of symptoms endorsed for each participant. For these regression calculations (3) individual age, gender, and scanner type were accounted for as covariates. Structural volume estimates of the LC were normalized by TBV, then also regressed onto the attention scores (4). Regression analyses were also replicated with handedness data included.

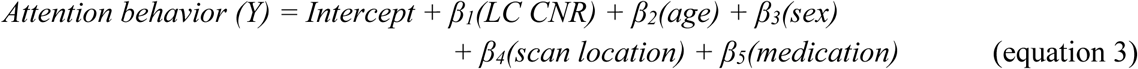

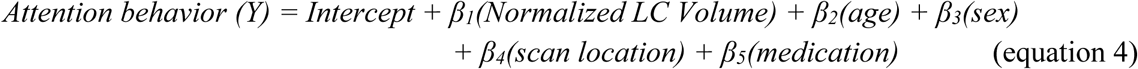

## Mediation analysis

Given results implying inverse relationships between LC CNR to inattention and LC volume to inattention, we tested mediation using the lavaan SEM package within JASP 0.18.3 (Love et al., 2019; Rosseel, 2012). Our main interest of the model was to test whether right LC CNR was associated with inattention score through right LC normalized volume. The magnitude and the significance of the effect was calculated using 5000 bootstrapping resampling and a bias-corrected confidence interval. A confounder adjustment for age, gender, scan site and medication was performed on the mediation model.

## Results

### LC Neuromelanin contrast associated with attention scores

In examining the relationship between neuromelanin asymmetry and total score, we observed significant overall effects of CNR asymmetry in models for the total score and the subscales. As a result, we found that neuromelanin asymmetry was positively associated with the total attention score (Figure 2) and with the sub scores for hyperactivity and impulsiveness (Table 3). However, it was not significantly associated with inattentiveness.

**Figure 2.**
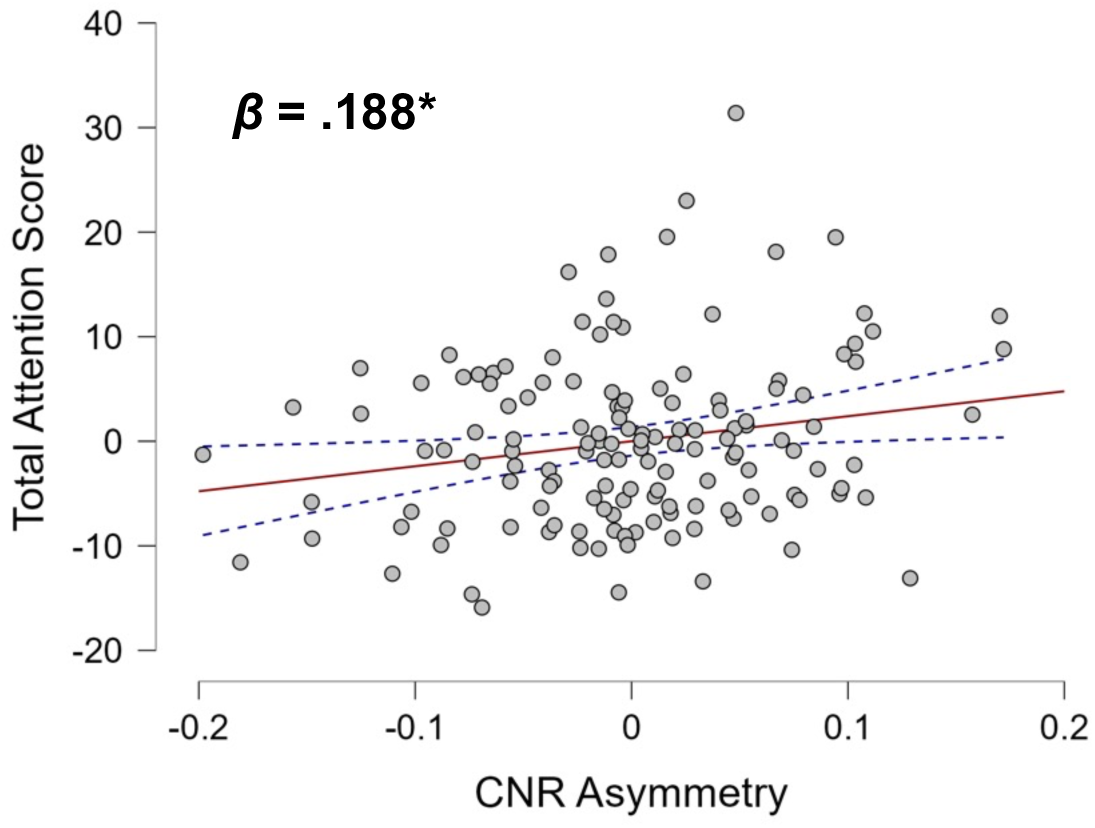
Partial regression plot of neuromelanin asymmetry onto total attention score Note. The regression line for neuromelanin asymmetry onto total attention score (red) is shown across residuals, in addition to 95% confidence interval (light blue). * p < .05

**Table 3.** Neuromelanin contrast asymmetry regressed onto attention scores.

| Behavior score | $\beta$ | p-value |
| --- | --- | --- |
| Total Attention Score | .188 | .019* |
| Inattentiveness | .125 | .127 |
| Hyperactivity | .221 | .006* |
| Impulsivity | .149 | .073† |

† $p < .10$ ; \* $p < .05$ ; \*\* $p < .005$

Follow-up regression analyses for each hemisphere of the LC, the left LC CNR fails to individually predict any of subscores or the total score, whereas the right LC CNR negatively predictive of inattentive behaviors (Figure 3 and Table 4). An additional t-test found that the left LC CNR (*M* =.181, *SD* = .064) is significantly higher than the right LC CNR (*M* = .129, *SD* = .059), *t* (140) = 8.868, p <.001.

**Figure 3.**
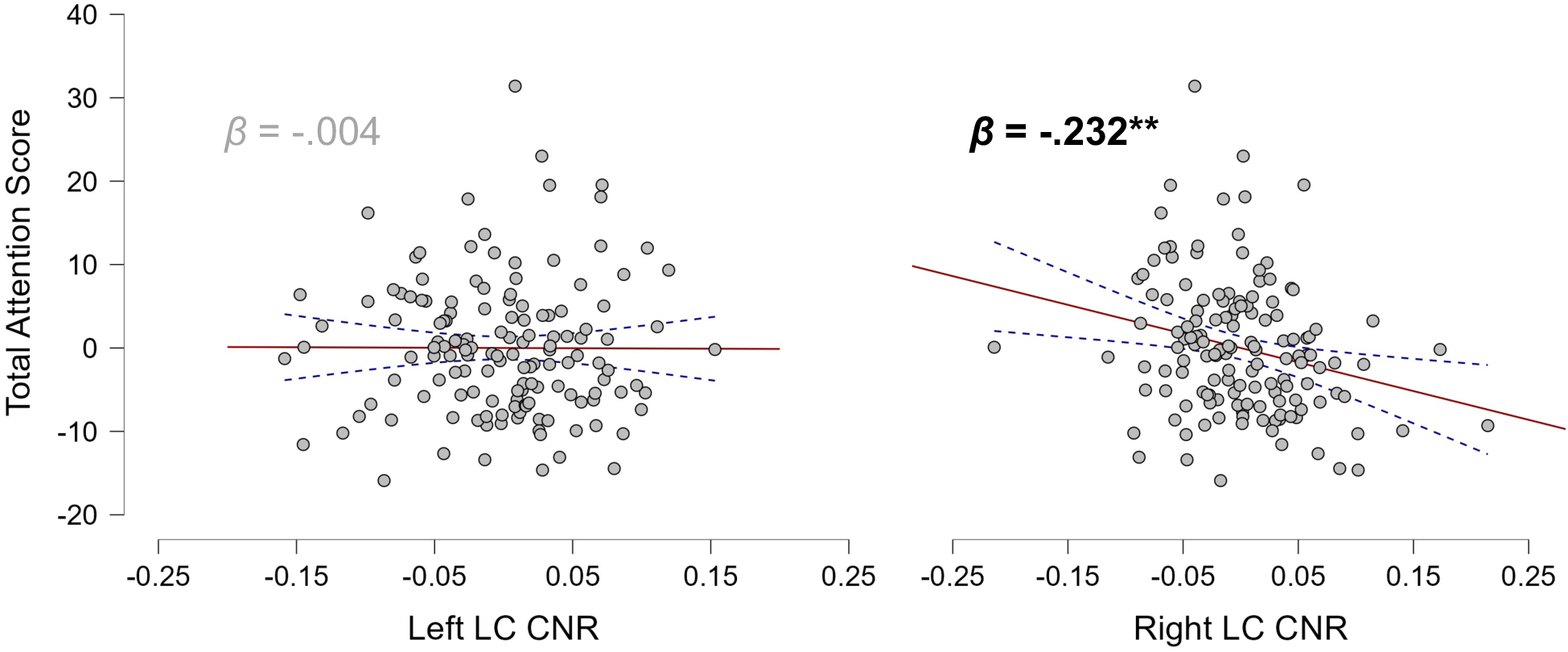
Partial regression plots of neuromelanin contrast by hemisphere onto total attention score, **p < .005

**Table 4.** Hemispheric neuromelanin CNR regressed onto behavior scores.

| Behavior score | LC CNR predictor | $\beta$ | p-value |
| --- | --- | --- | --- |
| Total Attention Score | Left hemisphere | -.004 | .961 |
|  | Right hemisphere | -.232 | .004** |
| Inattentiveness | Left hemisphere | -.053 | .544 |
|  | Right hemisphere | -.204 | .014* |
| Hyperactivity | Left hemisphere | .112 | .197 |
|  | Right hemisphere | -.157 | .057† |
| Impulsivity | Left hemisphere | -.032 | .723 |
|  | Right hemisphere | -.212 | .012* |

† $p < .10$ ; \* $p < .05$ ; \*\* $p < .005$

## LC Volume Estimation associated with attention scores

Hemispheric volume asymmetry did not predict the total score or sub-scores, including hyperactivity or impulsivity, but was found to be significantly predictive of inattentiveness (Table 5). Consistent with the LC CNR results, the left LC volume was not predictive of the total or inattentive scores, whereas the right LC volume showed a significant association with total score (β = .190, p =.024; Figure 4) and inattention (β = .282, p < .001). The right LC volume was not predictive of hyperactivity or impulsivity, but the left LC volume was positively predictive of hyperactivity behaviors (β = .179, p =.034). However, there was no volume difference by hemisphere (Left volume: *M* =45.411, *SD* = 12.112; right volume: *M* = 46.184, *SD* = 12.453; *p* = .565).

**Figure 4.**
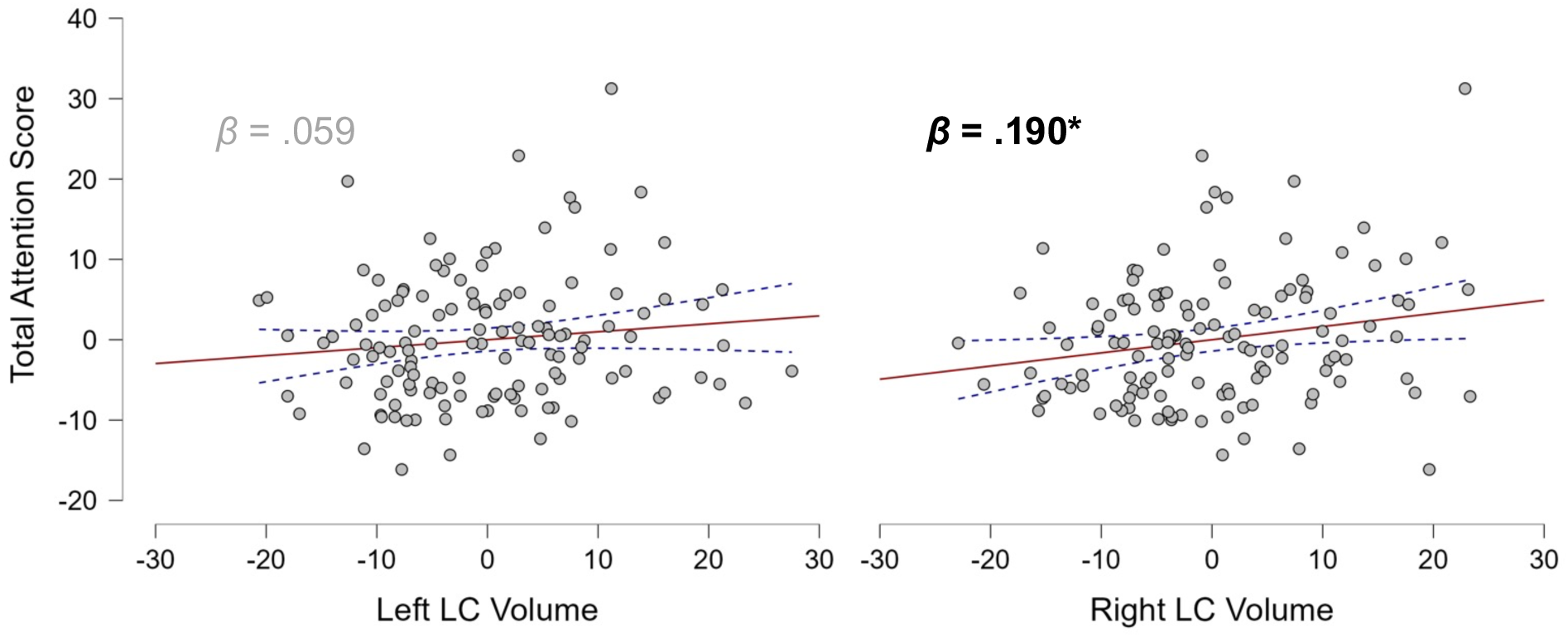
Partial regression of left and right LC volumes onto total attention score. *p < .05

**Table 5.** LC volumes regressed onto attention scores.

| Behavior score | LC volume predictor | $\beta$ | Predictor p-value |
| --- | --- | --- | --- |
| Total Attention Score | Asymmetry | -.059 | .482 |
|  | Left hemisphere | .112 | .179 |
|  | Right hemisphere | .190 | .024* |
| Inattentiveness | Asymmetry | -.171 | .044* |
|  | Left hemisphere | .059 | .487 |
|  | Right hemisphere | .282 | <.001*** |
| Hyperactivity | Asymmetry | .077 | .365 |
|  | Left hemisphere | .179 | .034* |
|  | Right hemisphere | .081 | .351 |
| Impulsivity | Asymmetry | .061 | .481 |
|  | Left hemisphere | .063 | .472 |
|  | Right hemisphere | -.017 | .852 |

\* $p < .05$ ; \*\* $p < .005$ ; \*\*\* $p < .001$

## LC Neuromelanin-Volume Interaction

Model 1 showed a significant indirect effect of LC CNR on inattention through normalized LC volume (indirect effect: β = −.064, SE = .031, p < 0.05, 95% CI [−.160 − -.013]; direct effect: β = −.140, SE = .081, p = .083, 95% CI = [−.301 −.025], Fig. 5). To rule out the possibility that LC volume’s association with inattention is mediated by neuromelanin, we also tested another model using LC volume as the independent variable, inattention score as outcome and LC CNR as mediator. This second mediation model was not statistically significant (indirect effect; p = .132, 95% CI = [.002 0.091]).

**Figure 5.**
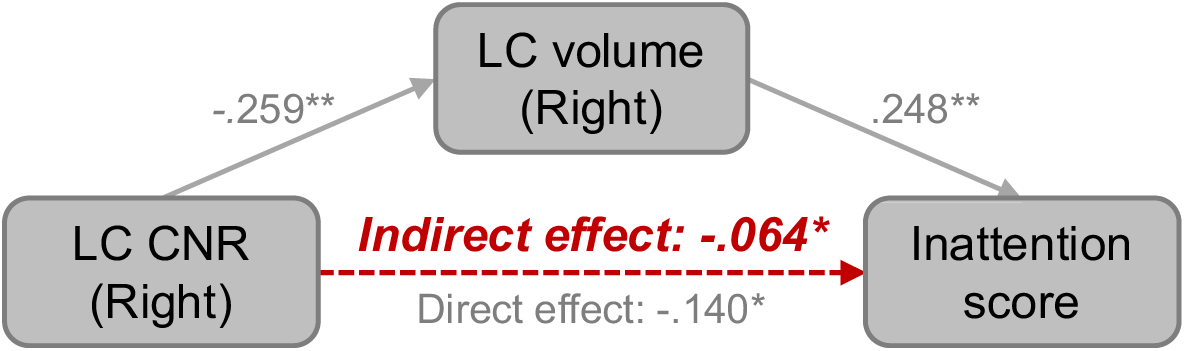
Mediation model of right neuromelanin contrast, right LC volume, and inattention score (*p < .05; **p < .005)

## Discussion

The present findings suggest a significant association between neuromelanin and attention behavior within a non-clinical, non-geriatric population. This is a notable expansion for the potential use of neuromelanin imaging to investigate the relation between LC and attentional behaviors in human populations, as prior to this study there have been concerns that gradual neuromelanin deposition patterns would make sensitive contrasts difficult to use beyond older adult populations (Galgani et al., 2023). In addition, as discussed previously, volume estimates of the LC have been difficult to perform due to the relatively small size of the region and proximity to respiratory and circulatory transports. The average LC sizes of 45-46 mm^3^ per hemisphere are in line with other multi-atlas estimates, and appear reasonably close to histological estimates, considering the relatively wide age range of the sample.

In addition to the LC’s relationship with inattentiveness, limited relationships between LC values and other behavioral patterns (hyperactivity and impulsivity) were observed. Neither set of scores was clearly associated with a hemisphere or valuation of the LC across CNR and volume, but there are reasons to believe this may be due to the cross-sectional nature of the sample. The latent presentation of hyperactivity has been observed to decline in adulthood, relative to other related behaviors (Martel et al., 2012), and model structures for adults have been best fit when accounting for differences in motor versus verbal hyperactive/impulsive behaviors (Gibbins et al., 2012; Gomez & Stavropoulos, 2021). This suggests that measures for these behavior sets may need to be reconceptualized for study in adults.

One specific finding that stands out in terms of how each predicts inattentive symptoms is the inverse relation of neuromelanin contrast to LC volume. The neuronal density of the LC has been identified as highest in the center (German et al., 1988), identical to the intensity of neuromelanin deposits as measured by contrasts. In neurodegenerative contexts decline in neuromelanin contrast is found to be associated with symptom progression, as is reduced volume in the substantia nigra, a neuromelanin-sensitive region more pliable for volume estimation (Biondetti et al., 2020; Kashihara et al., 2011; Li et al., 2022). In terms of mechanisms, disease progression is understood to lead to neuronal death, which reduces the overall volume and releases neuromelanin from within the neurons, with the neuromelanin then dissipating.

Regarding non-neurodegenerative contexts, in a comparative study the contrast ratios of the substantia nigra in schizophrenia patients were found to be higher than healthy controls, and the contrast ratio for the LC in depressive patients was found to be significantly lower than control (Shibata et al., 2008). A separate examination for noradrenergic dysfunction in cocaine users (W. Wang et al., 2021) found significantly higher LC contrast ratios amongst users compared to controls. In combination with functional deactivations across other regions, this was believed to be part of a neurotoxic effect of cocaine on the LC-NA system. Different neurons located throughout the LC have been associated with varying composition and projections targets (L. A. Schwarz & Luo, 2015). A larger than average LC may then consist of an abnormally distributed set of noradrenergic neurons, with reduced signaling to frontal and prefrontal regions, and increased norepinephrine transmission to other brain regions.

Beyond establishing basic associations between neuromelanin deposits and broad psychological symptoms, future analyses should aim to more effectively discern the implications of neuromelanin accumulation across the lifespan. Though metabolic connections between neuromelanin deposits and dopaminergic and norepinephrine producing neurons has been understood for some time (see Nagatsu et al., 2023), the precise mechanistic process still appears uncertain (Haining & Achat-Mendes, 2017; Priovoulos et al., 2020). Given that present theories as to this process are largely defined in the neurodegenerative context (Zucca et al., 2017), study of neuromelanin accumulation separately across the lifespan appears vital to properly interpret the molecule’s presence in alternative contexts. Examination of neuromelanin accumulation in the substantia nigra identified discrete periods of neuromelanin accumulation prior to adulthood (Fedorow et al., 2006), and further study for this possibility in both the nigra and the LC may be vital for understanding neurodevelopment.

Another finding of the analyses worth further discussion is the extent to which LC structure and metabolic byproduct were found to be lateralized by hemisphere. Previous investigation of psychopathology and the LC have observed similar patterns (Kumano et al., 2022; Llorca-Torralba et al., 2022; Tramonti Fantozzi et al., 2021). Lateralization of differences has also been observed in cortical regions as part of pathology (Levitt et al., 2002; Lindell, 2020; Oertel et al., 2010), in addition to longstanding observations of lateralized cognitive functions (Agcaoglu et al., 2022; Desmond et al., 1995; Holland et al., 2007; Spielberg et al., 2011). Lateralized attention processing has been previously observed within the right hemisphere (Bartolomeo & Seidel Malkinson, 2019; Mengotti et al., 2020), but has been contextualized primarily within dorsal and ventral attention networks to date. Examination of right hemisphere functional connectivity within integrative networks such as the FPN or default-mode network (DMN) may discern lateralization patterns in executive processing as part of attention. The lateralization of metabolic adjustments in the right hemisphere of the LC being associated with inattentive symptoms should be noted, as these provide insight both into cognitive connections to the LC and a more specific region of interest for possible use as a biomarker.

There may be a number of challenges in modeling connections between LC volume or neuromelanin and cortical networks in the context of inattention. The small size of the LC has been known to make size estimation difficult, and therefore LC analyses struggle to evaluate the effects of structure. As a metabolic byproduct of norepinephrine production, neuromelanin may best be understood as a marker related to broad function for the LC. This may not fully capture the relation between specific neural connections of the LC to cortical networks, which may be better observed through using neuromelanin sensitive images to improve functional connectivity analysis (Mäki-Marttunen & Espeseth, 2021; Turker et al., 2021).

Though neuromelanin-sensitive MRI contrasts represent a substantial, if uncertain, marker of LC norepinephrine production, resting state functional connectivity can provide more direct evidence of specific neural abnormalities. The differentiations can be both by specific neural region (Huang et al., 2021), or by abnormality in signal pattern (Gong et al., 2021). In the context of feedback sensitive tasks such as selective attention, measures such as dynamic functional connectivity may represent an intrinsic LC function associable with specific cognitive actions and impairments (Damaraju et al., 2014; Fong et al., 2019; Hutchison et al., 2013). While traditionally challenged by the small size of the LC, emerging techniques such as neuromelanin contrast seeding and 7T imaging allow for improved functional connectivity analysis of the LC, such as the ability to discern differential patterns across the LC itself (Betts et al., 2017). Differential effects of LC functional connectivity have also been observed previously across the lifespan (Jacobs et al., 2018; Song et al., 2021), and expanding upon such temporal findings will further improve the context into which LC connectivity patterns are placed.

In addition to structural alteration being associated with psychopathology, the functional connectivity of the LC may also significantly differ in the prevalence of disorders. Decreased LC functional connectivity to sensorimotor networks in ASD individuals has been associated with motor deficits and symptoms (Y. Huang et al., 2021). In individuals who have experienced trauma, LC signaling may mediate the prevalence of hyperresponsivity symptoms (Naegeli et al., 2018). Within older adults abnormal LC connectivity with the FPN has been associated with increased inattentive symptoms (Lee et al., 2018). In terms of normative patterns, LC functional connectivity has been found to exhibit a curvilinear pattern, increasing through adolescence and young adulthood, and declining as part of typical cognitive changes in older adults (Jacobs et al., 2018; Song et al., 2021). Song et al.’s study found that LC connectivity with sensory regions is high in early childhood and older adults but low with frontal regions such as frontal pole and the frontal medial cortex. The inverted curves of functional connectivity with the LC coincide with reduced attention performance for the respective age groups, indicating sensory overflow is prevalent for both (hypersensitivity) but attentional control is either underdeveloped in children, or has suffered degeneration in older adults.

Recently, studies suggested that a potential pathway for dysfunction through these regions and networks may include inconsistent/insufficient signaling between the LC and the SAL. This may operate through a form of bottom-up signaling, in which LC abnormalities would influence the SAL, which in turn limits the ability of the SAL to either activate the FPN/DMN or deactivate the opposing network in response to appropriate stimulus (Lee et al., 2020). On the other hand, a top-down model of neuro-regulation suggests that dysfunctions from cortical networks such as SAL and FPN could signal abnormally to the LC, affecting its performance (Unsworth & Robison, 2017). These two opposite models of LC-SAL direction are not necessarily mutually exclusive, as neural circuits may seek both up and down regulation at different points (Aston-Jones & Cohen, 2005). While directionality in the relationship may be a point of continuing study, present findings support the importance of the LC-SAL connection. A further benefit of such a model is that it provides insights into the mechanisms linking connections to impulsivity and clinical conditions such as ADHD, by virtue of the SAL’s connections to impulse control. The SAL, through both structural deficits (Galandra et al., 2018; Grodin et al., 2017) and limited between-network connectivity (Hobkirk et al., 2019; Navalpotro-Gomez et al., 2020) has been associated with increased impulsivity. Parsimonious models of dysfunction within those with ADHD could benefit from the network’s association with multiple symptom clusters.

The distinct possibility that variance in LC signaling may be responsible for cortical network differences is compelling due to potentially resolving conflicting findings and by providing a common origin across several disorders. Though the evidence for a signaling pathway from the LC through the SAL and into frontal networks for attentional processes is compelling, it should be noted that alternative models of attention exist, such as those considering feedback. Unsworth and Robinson (2017) posited a model including the LC-NA system with specific signaling connectivity to cortical networks, but with frontal networks performing positive feedback signaling through the SAL to the LC. At the same time, findings such as DMN-FPN anti-correlation being positively associated with cognitive performance (Keller et al., 2015) and regulated LC-NA signaling associated with optimal function (Aston-Jones & Cohen, 2005) suggest each network or region is vital to the overall neural performance of this potential pathway. Thus variance in the function of one network may be causative of a plurality of neural attention correlation, but not comprehensive in explaining performance. The possibility of heterogeneous dysfunctions within an attention networking model would appear supported by the diversity of symptom presentations in any given individual.

One additional aspect of a mechanistic neural model worth exploring is a possible identification of structural differences related to the function of previously identified neural regions. Though frequently studied as separate modalities, structural variances have been previously associated with functional connectivity, including large-scale networks and relations to psychopathology (de Kwaasteniet et al., 2013; Greicius et al., 2009; Korponay et al., 2017;

Marstaller et al., 2015). A recent study observed functional relationships between the LC, cortical networks, and attention performance (Neal et al., 2023). More specifically, the findings there included significant positive associations between the LC functional connectivity to SAL regions (including insula and dorsal ACC) and attention performance. The same analyses also observed an association between SAL functional connectivity to the right DLPFC and attention performance, further supporting previous lateralized network findings. Given previous work establishing connectivity between the LC and cortical network regions linked to attention, evaluating the possibility of structural differences that may underlie or augment connectivity differences would assist in developing a more holistic model.

Beyond methodological refinement of neuroimaging components, future studies can and should attempt to place the observed relations here within a more robust clinical context. Previous literature regarding ADHD, a disorder of particular focus due to the centrality of neurodevelopmentally-related inattention to presentation, has observed relations between these symptoms and alterations in frontal structure or function (Almeida et al., 2010; Baving et al., 1999; Loo et al., 2009). The alterations within the frontal region in particular have been associated with deficits in executive control and attentional task performance (S. Wang et al., 2013). Thus, the degree of structural and functional alterations in the frontal region may be useful for discriminant, individual identification of those with ADHD (Cheng et al., 2012; Iannaccone et al., 2015). Given previously described interactions between cortical and subcortical regions, neuroplasticity within these circuits may mean that initial abnormalities within one region may shape errant development within a corresponding portion of the brain. A prenatal or neonatal condition affecting the LC’s capacity to sufficiently signal to cortical, attention-associated regions may lead to reduced connectivity and preserved structure across development, to suggest one possibility.

The frontal cortex abnormalities are not the only observed neurological difference for attention deficits and ADHD, however. Lower levels of cortical thickness have been observed for those with the disorder (Narr et al., 2009; Silk et al., 2016). A meta-analysis of subcortical structural differences between ADHD and TD individuals found reduced volumes for those with ADHD in a number of subcortical regions, including the nucleus accumbens, amygdala, caudate, hippocampus, and putamen, as well as reduced overall intracranial volume (Hoogman et al., 2017). Again, the potential for midbrain projections from the LC to be altered in a way that disrupts norepinephrine transmission within these midbrain regions may impact structural composition across development. The functional connectivity of subcortical regions has also been identified as substantially different in those with ADHD (Costa Dias et al., 2013; Kowalczyk et al., 2022; E. M. Miller et al., 2012). Additional analyses of such projections and corresponding regions across developmental periods and within clinical populations may observe a more integrated set of findings than observed presently.

Noradrenergic pathways are not the only systems within subcortical imaging worth further study for possible origins of attention deficits and related psychopathology.

Dopamine is a neurotransmitter associated with neural reward incentives and impulse control (Dalley & Roiser, 2012), processes identified as aberrant alongside attention in ADHD.

Additionally it shares a biosynthetic pathway and similar cortical regions of signaling with norepinephrine (Ranjbar-Slamloo & Fazlali, 2020), suggesting examination of both may be valuable for a holistic understanding of differences in attention and related executive functions. Dopamine has been observed improving synchronization between resting state networks (Dang et al., 2012). A genetic connection to dopamine transport has been found related to abnormal striatal functional connectivity in children with ADHD (C.-Y. Shang et al., 2021). Abnormalities in microstructure associated with dopamine transmission in the striatum have been associated with Parkinson’s disease (S. Shang et al., 2021).

The substantia nigra and ventral tengmental area are two main regions of dopamine production within the brain, with differing cortical projections nonetheless similarly associated with reinforcement and inhibition (Ilango et al., 2014). These functions, as well as an established link between dopaminergic degradation and psychopathology in the neurodegenerative context (Alberico et al., 2015; Bae et al., 2021; Q. Chen et al., 2021; Parent & Parent, 2010), suggest that examination of these regions in the context of other disorders and symptoms may be valuable.

Given common metabolic pathways to dopamine and norepinephrine production and observed relationships to attention-related symptoms, a common catecholaminergic dysfunction has been suggested as a mechanism for neurodevelopmental disorders such as ADHD (Prince, 2008).

Serotonin is a neurotransmitter associated primarily with affect (Jauhar et al., 2023; Jones et al., 2020; Michely et al., 2020), as well as metabolic impulses such as food and sleep (van Galen et al., 2021; Vaseghi et al., 2022). Beyond the phenotypic connections between these effects and attention serotonergic and noradrenergic systems are known to interact (Blier, 2001; Guiard et al., 2008), a commonality treated by the use of serotonin and norepinephrine reuptake inhibitors across psychopathology (Dezfouli et al., 2024; Gabriel & Violato, 2011; Gorman, 2000; Hartford et al., 2007). For examination of regions specific to serotonin production the raphe nuclei has been previously identified as the main region within the brain (Beliveau et al., 2015; Hornung, 2003; Maier & Watkins, 2005). Examination of this region due to potential interactions with neural attention processes could be another step towards a holistic understanding of this aspect of cognition.

Acetylcholine is an additional neurotransmitter associated with sustained attention, as well as learning and memory. Cholinergic enhancers have been found to improve task performance and cortical activation in attention and emotion processing tasks (Bentley et al., 2003). White matter pathways connected to attention and memory performance in older adults (Nemy et al., 2020). Neural degradation in cholinergic producing regions has been associated with reduced attention and memory performance in Alzheimer’s (Machado et al., 2020; Nemy et al., 2023) and Parkinson’s patients (Crowley et al., 2024; Zee et al., 2021). Additionally common cholinergic and dopaminergic connections to the striatum have been observed with interaction between the two (Myslivecek, 2021), suggesting effects beyond basic associations with cholinergic effects alone. Neurodevelopmental abnormalities to these same symptoms may produce similar deficits in executive functioning abilities in younger populations.

Additionally, emerging understandings of neurotransmitter production and transmission have identified the possible influence of extracranial sources of influence such as gut microbiomes relating to symptomatology and neurodevelopment (Gkougka et al., 2022), emphasizing the further need for holistic study in this area.

ADHD research also encompasses a set of developmental theories related to neurological structure and function. One such conceptualization is the maturational delay theory, in which the core neurological dysfunction in ADHD is a set of structural and connective patterns which eventually develop, but are delayed compared to typical development. This pattern has been found in both structural and functional measure of individuals with the disorder (Shaw et al., 2007; Y. Wang et al., 2021). However, some individuals exhibit ADHD symptoms into adulthood, indicating that maturational delay cannot be explained by a single theoretical approach encompassing all individuals with the disorder. For adults with continued ADHD presentation, a high degree of structural similarity with child ADHD individuals was found (Zhang-James et al., 2021), though structural differences, compared to typical development, were noted as more discriminant in children. A resting-state fMRI study noted higher functional connectivity between the DMN and limbic network as discriminant of adults with ADHD versus children with the disorder (Guo et al., 2020) suggesting that more thorough examination of the different progressions of the disorder may also reveal different associations between abnormalities in structural regions or functional connectivity.

One possible method to best examine these possibilities would be a large-scale, longitudinal study examining structural and functional relationships across neurodevelopment. Extant projects, such as the ABCD study (Barch et al., 2018), provide the capacity to discern aspects of pathology such as continuous presentation from childhood or alternative mechanisms for the reported symptoms in combination with neuroimaging techniques represent an important expansion of the literature. Standardized inclusion of measures such as neuromelanin sensitive MRI in such batteries would assist in broader examination of neural relationships across pathologies and development.

These findings improve our knowledge of the subcortical substrates linked to inattentive symptoms suggested by previous findings within neurodegenerative structural and functional MRI. Inattentive symptom reporting was associated with multiple structural values in the LC. Neuromelanin contrasts were observed in association with inattention-related behaviors, and specific subscores were related to the left and right hemispheres of the LC, furthering details of the subregion’s functions. The association of specific LC dimensions to attention behaviors provides additional insight for holstic models of inattention, such as those consolidating other structural or functional differences throughout the brain.

## Acknowledgements

Grants and funding: This work was done based on research supported by R01AG075000/AG/NIA NIH and by 2022 Virginia Tech Lay Nam Chang Dean’s Discovery Fund.

## Notes

### Competing Interest Statement

The authors have declared no competing interest.

